# The lncRNA TRG-AS1 promotes the growth of colorectal cancer cells through the regulation of P2RY10/GNA13

**DOI:** 10.1101/2023.07.04.547664

**Authors:** Lin Zhuang, Baoyang Luo, Linghui Deng, Qi Zhang, Yuanjiu Li, Donglin Sun, Hua Zhang, Qiutao Zhang

**Affiliations:** The First People’s Hospital of Changzhou, The Third Affiliated Hospital of Soochow University, Changzhou, Jiangsu, China; Department of General Surgery, Wujin Affiliated Hospital of Jiangsu University and The Wujin Clinical college of Xuzhou Medical University, Changzhou, Jiangsu, China; Department of Hepatobiliary and Pancreatic Surgery,The Affiliated Taizhou People’s Hospital of Nanjing Medical University,Taizhou,Jiangsu,China; Department of Oncology, Wujin Affiliated Hospital of Jiangsu University and The Wujin Clinical college of Xuzhou Medical University, Changzhou, Jiangsu, China; Department of oncology, The First Affiliated Hospital of Soochow University, Suzhou, Jiangsu, China

**Keywords:** colorectal cancer, TRG-AS1, LoVo cells, P2RY10, GNA13

## Abstract

**Purpose:** The lncRNA TRG-AS1 and its co-expressed gene P2RY10 are important for colorectal cancer (CRC) occurrence and development. The purpose of our research was to explore the roles of TRG-AS1 and P2RY10 in CRC progression.

**Methods:** The abundance of TRG-AS1 and P2RY10 was determined in CRC cell lines. LoVo cells were transfected with si-TRG-AS1 and si-P2RY10 constructs. Subsequently, the viability, colony formation, and migration of the transfected cells were analyzed using cell counting kit-8, clonogenicity, and scratch-wound/Transwell® assays, respectively. Cells overexpressing GNA13 were used to further explore the relationship between TRG-AS1 and P2RY10 along with their downstream functions. Finally, nude mice were injected with different transfected cell types to observe tumor formation *in vivo*.

**Results:** TRG-AS1 and P2RY10 were significantly upregulated in HT-29 and LoVo compared to FHC cells. TRG-AS1 knockdown and P2RY10 silencing suppressed the viability, colony formation, and migration of LoVo cells. TRG-AS1 knockdown downregulated the expression of P2RY10, GNA12, and GNA13, while P2RY10 silencing downregulated the expression of TRG-AS1, GNA12, and GNA13. Additionally, GNA13 overexpression reversed the cell growth and gene expression changes in LoVo cells induced by TRG-AS1 knockdown or P2RY10 silencing. *In vivo* experiments revealed that CRC tumor growth was suppressed by TRG-AS1 knockdown and P2RY10 silencing.

**Conclusions:** TRG-AS1 knockdown repressed the growth of CRC cells by regulating P2RY10 and GNA13 expression, thereby controlling CRC occurrence and development.

## 1 Introduction

Colorectal cancer (CRC) is one of the most prevalent malignancy around the world and in 2020, there were an estimated 1,900,000 new cases and 900,000 deaths globally [1]. The incidence of CRC has continued to increase, currently accounting for approximately 10 percent of all cancers, and it is the second most frequent reason for cancer-related death [2]. Despite continuous technological development and increasing research into comprehensive CRC treatment, radical surgical excision, radiotherapy, targeted therapy, and chemotherapy remain the primary therapeutic methods [3]. Although drug treatment options continue to evolve, breakthroughs in metastatic CRC treatment are difficult to achieve, and the prognosis is still poor, with a median overall survival (OS) of only 25–30 months [4]. Immune checkpoint blockade (ICB) therapy enables some patients to achieve lasting benefits and significantly improve disease outcomes [5]. Programmed death-1 (PD-1)-targeting ICB therapies, such as pembrolizumab and navulizumab, have been approved for the treatment of CRC with dMMR or high MSI-H [6]. However, dMMR/MSH-H is rare (approximately 15% of CRC patients and 4% of metastatic CRC patients), and some patients rapidly develop immune resistance [7, 8]. Consequently, there is an urgent need to investigate the molecular mechanisms underlying CRC and to identify potential therapeutic targets for CRC management.

Long non-coding RNAs (lncRNAs, >200 nucleotides) do not encode proteins or peptides, but have essential functions in a variety of biological processes by regulating gene expression at the post-translational, translational, and transcriptional levels [9]. Recent reports have illustrated that dysregulated lncRNA profiles are widely involved in tumor pathogenesis, including epithelial-to-mesenchymal transition (EMT), cell proliferation, and tumor resistance [10-12]. Our previous study found 428 DE lncRNAs in CRC tissues relative to normal tissues, and analysis of the lncRNA profiles in CRC revealed that the lncRNA TRG-AS1 and P2RY10 were co-expressed [13]. The lncRNA TRG-AS1 has been reported to be an essential lncRNA involved in shaping the tumor microenvironment in squamous cell carcinoma of the head and neck [14]. Zhang et al. [15] showed that lncRNA TRG-AS1 expression was abundant in lung cancer, and its overexpression promoted the proliferation and invasion of lung cancer cells via the miR-224-5p/SMAD4 axis, thus representing an underlying target for lung cancer treatment. P2RY10 is a G protein-coupled receptor that is activated by adenosine and uridine. Wang et al. [16] found that P2RY10 might be crucial for the immune response and development of metastatic melanoma. However, the specific functions of TRG-AS1 and P2RY10 in CRC remain unknown.

In addition, high P2RY10 expression can upregulate the expression of GNA12 and GNA13, which are essential for tumor cell proliferation [17]. It has been reported that GNA12/13 activation can accelerate proliferation and tumor formation in small-cell lung carcinoma, ovarian cancer, and hepatocellular carcinoma cells [18]. Therefore, the current study first determined the levels of P2RY10 and lncRNA TRG-AS1 in normal and CRC cell lines, and then explored the functions of TRG-AS1 and P2RY10 in CRC development *in vitro* and *in vivo*. These results provide further insights into CRC progression and pathogenesis and highlight novel targets and pathways for CRC diagnosis and treatment.

## 2 Materials and methods

### 2.1 Cell acquisition and culture

Human colon normal epithelial cells (FHC) and CRC cell lines (HT29 and LoVo) were obtained from Cell Bank, Chinese Academy of Sciences (Shanghai, China). Among them, both the FHC and HT-29 cells were cultured in Dulbecco’s modified Eagle’s medium (DMEM, Thermo Fisher Scientific) supplemented with 10% fetal bovine serum (FBS, Thermo Fisher Scientific) and 1% penicillin/streptomycin (Thermo Fisher Scientific); LoVo cells were cultured in Ham’s F-12K medium (Servicebio) containing 10% FBS and 1% penicillin/streptomycin. A 37 °C incubator with 5% CO_2_ was used to maintain all cells.

### 2.2 Real-time quantitative PCR (RT-qPCR)

Various cell lines were treated with RNAiso Plus (Takara, Beijing, China) to extract total RNA, based on the manufacturer’s recommendations, and then the isolated RNA (1 μg) was reverse transcribed into cDNA (20 μL) using PrimeScript™ RT Master Mix (Perfect Real Time, Takara). The qPCR reaction was initiated at 95 °C for 3 min, followed by 40 cycles at 95 °C for 10 s and 60 °C for 30 s. The sequences of all the primers used are presented in **Table 1**, with *GAPDH* as the reference gene. The 2^-ΔΔCT^ method was used to calculate the relative mRNA expression levels of lncRNA TRG-AS1, *P2RY10, GNA12*, and *GNA13*.

### 2.3 Cell transfection

Shanghai Generay Biotechnology (Shanghai, China) prepared and synthesized the si-lncRNA TRG-AS1-1/2/3, si-P2RY10-1/2/3, and si-negative control (NC). The negative control (NC) and GNA13 overexpression (OE-GNA13) plasmids were obtained from Sangon Biotech (Shanghai, China). Cell transfection was performed as previously described [19]. Briefly, LoVo cells (4 × 10^4^ cells/well) were inoculated into a 24-well plate and transfected with (1) si-lncRNA TRG-AS1-1/2/3 or si-NC, (2) si-P2RY10-1/2/3 or si-NC, and (3) OE-GNA13 or NC plasmids using Lipofectamine™ 2000 (Thermo Fisher Scientific) following the manufacturer’s recommendations. After 6 h of transfection, complete medium was supplemented, and the cells were cultured for another 48 h. LncRNA TRG-AS1, *P2RY10*, and *GNA13* expression was detected using RT-qPCR to evaluate cell transfection efficiency.

### 2.4 Cell viability and colony formation assays

LoVo cells transfected with different plasmids (1 × 10^4^ cells/well) were seeded into a 96-well plates. After culturing for 24, 48, and 72 h, 10 μL of cell counting kit-8 (CCK-8, Beyotime Biotechnology, Shanghai, China) reagent was added to each well, and incubated for another 2 h. A microplate reader (Thermo Fisher Scientific) was used to measure the absorbance values at 450 nm.

For colony formation, the transfected LoVo cells were harvested and resuspended in 3 mL of complete medium (adjusted to 100 cells/mL). Next, 500 cells were added to each well and incubated for 14 d. After the cell colonies became visible to the naked eye, they were washed twice with phosphate buffered saline (PBS) and fixed with 4% paraformaldehyde at room temperature for 10 min. After removing the fixative, the cells were stained with crystal violet for 10 min, washed, dried, and imaged.

### 2.5 Cell migration assays

Scratch-wound and Transwell® assays were used to monitor cell migration. For the scratch-wound assay, horizontal lines (approximately 1 cm) were drawn on the back of a 6-well plate, and different cell samples were seeded in the plate. On the second day, a scratch was made perpendicular to the horizontal lines on the plate and serum-free medium was added to each well. Cells were imaged after 0 and 48 h of culture.

For the Transwell® assay, the transfected LoVo cells were harvested and resuspended (2 × 10^5^ cells/mL). A total of 200 μL of cell suspension was added to the upper Transwell® chambers (pore size 8 μm; Guangzhou Jet Bio-filtration Co., Ltd., Guangzhou, China) in a 24-well plate, and the lower chamber was filled with complete medium (500 μL). After 48 h, the Transwell® chambers were taken out, the cells were collected, fixed and stained for 20 min. After removing the excess dye, the cells were imaged using an inverted microscope.

### 2.6 Western blot

LoVo cells with different plasmid transfections were lysed with 200 μL RIPA lysis buffer, and total protein was isolated from the cells. A BCA protein assay kit (Beyotime Biotechnology) was used to determine protein concentration. Next, the protein samples (20 μg) were separated using 10% SDS-PAGE and transferred to PVDF membranes. The membranes were blocked with 5% skim milk at 37 °C for 2 h, then washed three times with 1X PBST (1000 mL 1X PBS + 1 mL Tween-20), and incubated with anti-P2RY10, anti-GNA12, anti-GNA13, or anti-GAPDH antibodies at 4 °C overnight. Thereafter, membranes were washed and incubated with secondary antibodies (1:10,000, Jackson ImmunoResearch Laboratories, Inc.) at 37 °C for 2 h. After washing five times with PBST, the protein bands were visualized using an automatic chemiluminescence image analysis system (Tanon Science & Technology, Shanghai, China), and quantified using ImageJ software (version 1.8.0, USA).

### 2.7 Tumor formation in nude mice

Cells subjected to different plasmid transfections (si-NC, si-TRG-AS1, or si-P2RY10) were harvested at the logarithmic growth stage and resuspended. PBS was used to adjust the cell density to 5 × 10^7^ cells/mL. Nine female BALB/c nude mice (6–8 weeks old) were purchased from Shanghai SLAC Laboratory Animal Co., Ltd. (Shanghai, China) and maintained under specific pathogen-free conditions. After a week of adjustable feeding, all nude mice were randomly divided into three groups: si-NC, si-TRG-AS1, and si-P2RY10 (n = 3 per group). The mice were subcutaneously injected with 100 μL of si-NC, si-TRG-AS1, or si-P2RY10 transfected LoVo cell suspension at the axillary region of the right forelimb. All mice were monitored daily, and tumor volumes were determined weekly. After five weeks of induction, all mice were sacrificed by cervical dislocation, and the tumors of each mouse were harvested. After measuring the tumor dimensions, tumor tissues from each mouse were fixed and stained with hematoxylin and eosin (H & E; Beyotime Biotechnology).

### 2.8 Statistical analysis

Data were shown as mean ± standard deviation from three independent experiments. GraphPad Prism 5.0 software (GraphPad Software, CA, USA) was used for statistical analysis. The difference between two groups was calculated using Student’s *t*-test, while one-way analysis of variance (ANOVA) with a post-hoc Tukey test was used to compare more than two groups. *P* < 0.05 was considered the threshold for statistical significance.

## 3 Results

### 3.1 Cellular expression levels of lncRNA TRG-AS1 and P2RY10

Our previous research indicated the dysregulation of TRG-AS1 and P2RY10 in CRC through bioinformatic analysis; therefore, to validate the previous observation, we measured TRG-AS1 and P2RY10 levels in CRC cells. The TRG-AS1 and P2RY10 expression levels were considerably higher (*P* < 0.05) in CRC HT-29 and LoVo cells than in FHC control cells (**Figure 1A**). The difference in expression levels between FHC and LoVo cells (*P* < 0.01) was more significant than that between FHC and HT-29 cells (*P* < 0.05). Therefore, LoVo cells were selected for the subsequent experiments.

**Figure 1.**
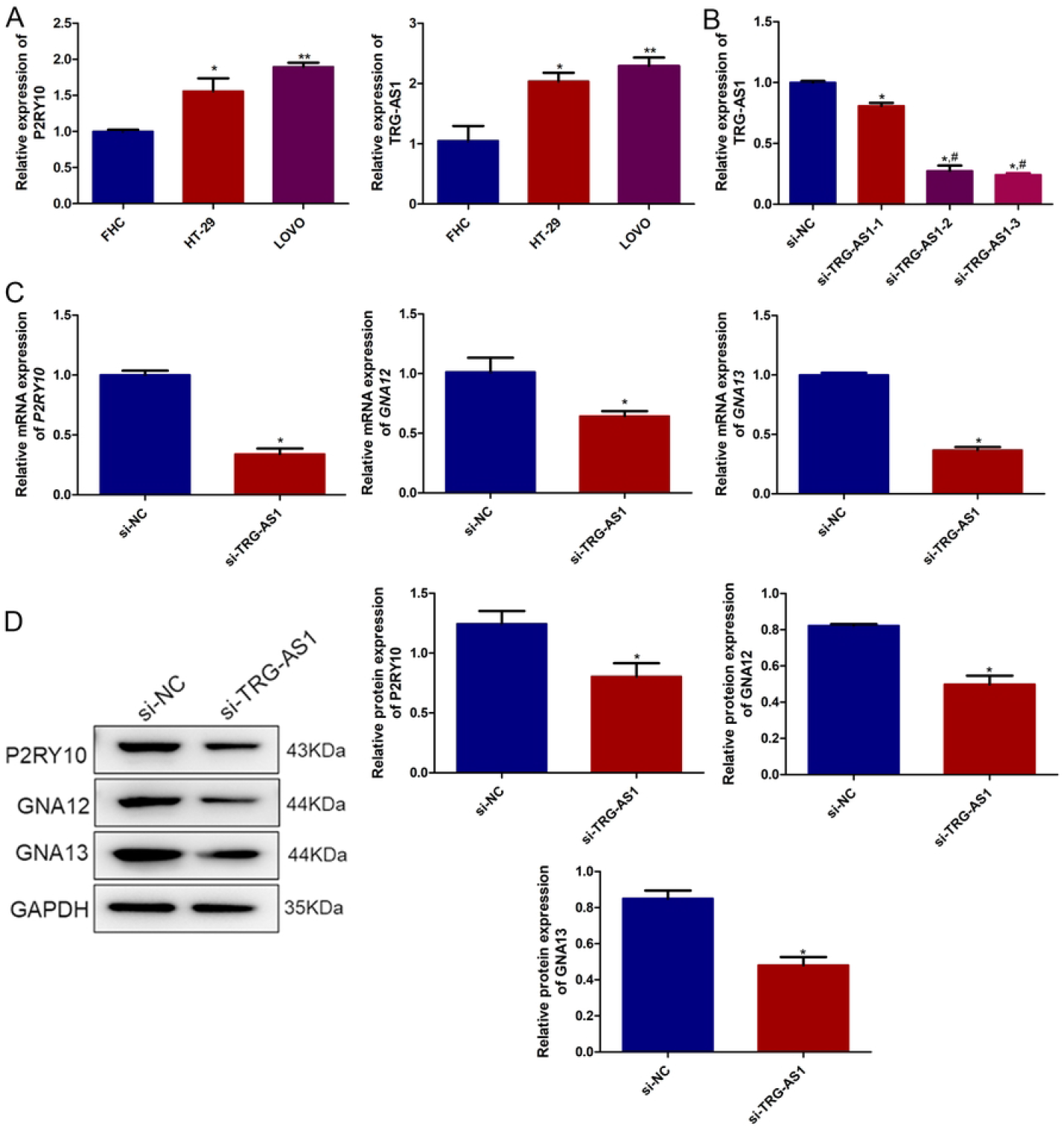
Effect of TRG-AS1 on the expression of P2RY10, GNA12, and GNA13 in LoVo cells. (**A**) Expression of P2RY10 and TRG-AS1 in human normal colon epithelial (FHC) and colorectal cancer (CRC) HT29 and LoVo cell lines. (**B**) Cell transfection efficiency of LoVo cells with si-TRG-AS1. (**C**) Expression of *P2RY10, GNA12*, and *GNA13* mRNA in transfected LoVo cells determined by RT-qPCR. (**D**) Western blot of P2RY10, GNA12, and GNA13 protein expression in transfected LoVo cells. * P < 0.05, ** P < 0.01 vs. FHC cells or si-NC; ^#^ P < 0.05 vs. si-TRG-AS1-1.

### 3.2 Effect of TRG-AS1 on P2RY10, GNA12, and GNA13 expression in LoVo cells

To investigate the role of TRG-AS1 in CRC, LoVo cells with TRG-AS1 knockdown were established by transfection of targeting siRNAs. Compared to the si-NC cells, TRG-AS1 levels in the LoVo cells transfected with si-TRG-AS1-1/2/3 were significantly decreased (*P* < 0.05). Cells transfected with si-TRG-AS1-2/3 were more significantly affected (*P* < 0.05) than the cells transfected with si-TRG-AS1-1 (**Figure 1B**). Therefore, si-TRG-AS1-3 was selected and successfully used to establish TRG-AS1 knockdown LoVo cells.

After TRG-AS1 knockdown, the mRNA and protein expression levels of P2RY10, GNA12, and GNA13 were determined using RT-qPCR and western blotting, respectively. Compared to si-NC cells, the relative mRNA and protein expression of *P2RY10, GNA12*, and *GNA13* was markedly downregulated (*P* < 0.05) in si-TRG-AS1-transfected LoVo cells (**Figure 1C, D**).

### 3.3 Role of TRG-AS1 in LoVo cell proliferation and migration

The effect of TRG-AS1 knockdown on the viability, colony formation, and migration of LoVo cells was assessed. After culturing for 24 h, the viability of LoVo cells transfected with si-NC or si-TRG-AS1 did not differ significantly (*P* > 0.05; **Figure 2A**). However, after culturing for 48 and 72 h, si-TRG-AS1 transfection significantly inhibited (*P* < 0.05) the viability of LoVo cells compared to si-NC cells (**Figure 2A**). LoVo cells transfected with si-NC or si-TRG-AS1 generated 453 ± 59.2 and 115.3 ± 99.5 colonies, respectively, indicating that si-TRG-AS1 had a pronounced effect on colony formation (*P* < 0.05; **Figure 2B**). Finally, Transwell® and scratch-wound assays were used to determine cell migration ability. The Transwell® assay results revealed that the number of transmigrated si-TRG-AS1 cells was evidently reduced compared to the si-NC cells (*P* < 0.05; **Figure 2C**). Similarly, the scratch-wound assay also showed that si-TRG-AS1 transfection significantly suppressed LoVo cell migration (**Figure 2D**). Taken together, these results indicated that TRG-AS1 knockdown significantly suppressed the viability, colony formation, and migration of LoVo cells.

**Figure 2.**
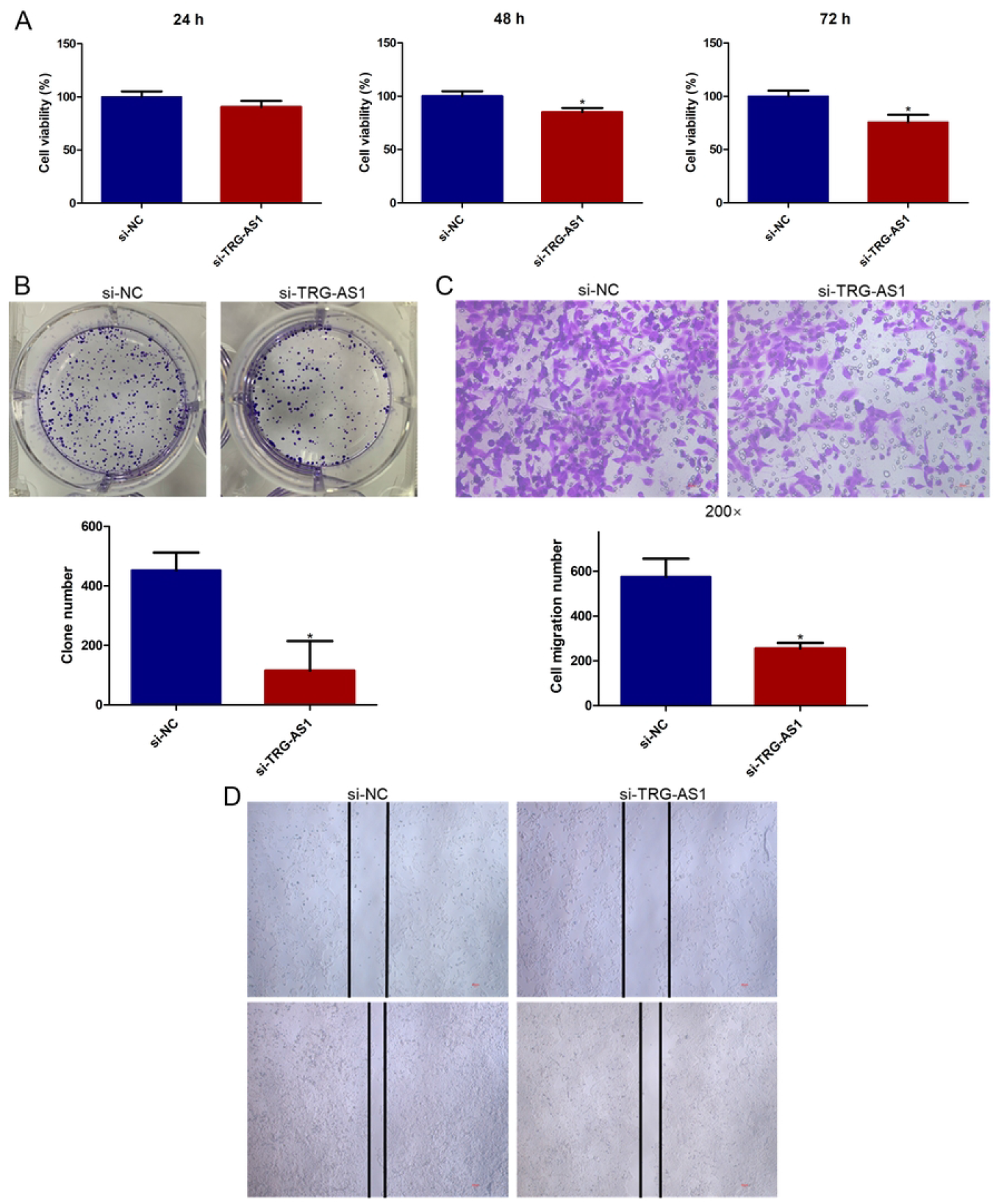
Effect of TRG-AS1 on LoVo cell proliferation. (**A**) Viability of LoVo cells after transfection with si-TRG-AS1 and cultured for 24, 48, and 72 h measured using the cell counting kit-8. (**B**) Transfected LoVo cell colony numbers by colony formation assay. Migration of transfected LoVo cells based on (**C**) Transwell® assay and (**D**) scratch test. * P < 0.05 vs. si-NC.

### 3.4 Effect of P2RY10 in TRG-AS1, GNA12, and GNA13 expression in LoVo cells

Our previous bioinformatics study found that P2RY10 is co-expressed with the lncRNA TRG-AS1 and is important for immune response. Owing to the increased level of *P2RY10* in LoVo cells, we generated LoVo cells expressing P2RY10 silencing constructs. Compared to si-NC cells, *P2RY10* expression was markedly downregulated in si-P2RY10-1/2/3 transfected cells (*P* < 0.05). Cells transfected with si-P2RY10-2/3 were more significantly affected (*P* < 0.05) than the cells transfected with si-P2RY10-1 (**Figure 3A**). These results indicated that P2RY10 was successfully silenced in LoVo cells. RT-qPCR and western blotting were used to determine the mRNA and protein expression of TRG-AS1, GNA12, and GNA13. Compared with si-NC cells, the mRNA expression of TRG-AS1, GNA12, and GNA13 was markedly downregulated in si-P2RY10 cells (*P* < 0.05; **Figure 3B**). Additionally, P2RY10 silencing markedly (*P* < 0.05) reduced P2RY10, GNA12, and GNA13 protein levels (**Figure 3C**).

**Figure 3.**
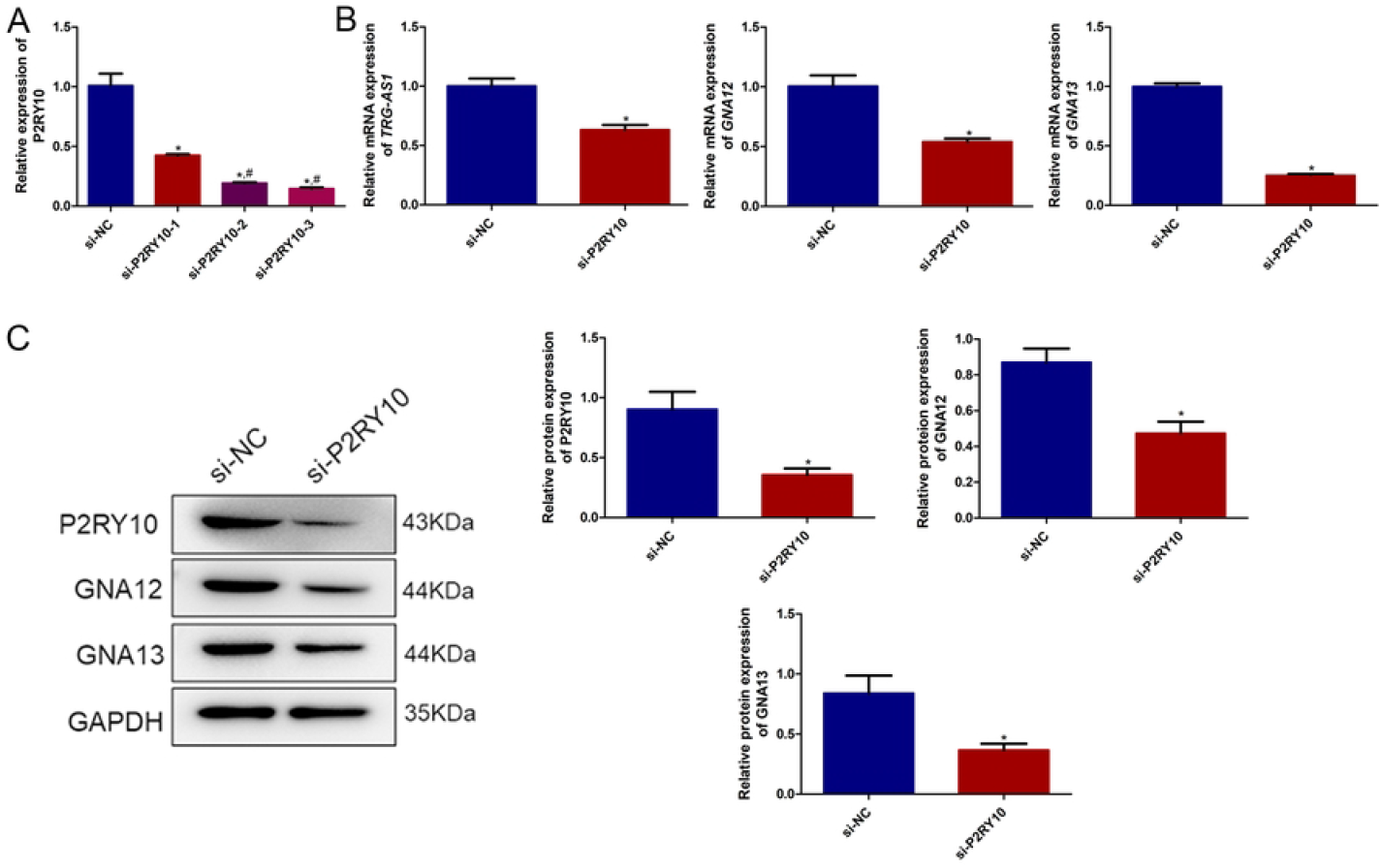
Effects of P2RY10 on the expression of TRG-AS1, GNA12, and GNA13 in LoVo cells. (**A**) LoVo cell transfection efficiency with si-P2RY10. (**B**) Expression of *TRG-AS1, GNA12*, and *GNA13* mRNA in transfected LoVo cells determined by RT-qPCR. (**C**) Western blot of P2RY10, GNA12, and GNA13 protein expression in transfected LoVo cells. * P < 0.05 vs. si-NC; ^#^ P < 0.05 vs. si-P2RY10-1.

### 3.5 Role of P2RY10 in the proliferation of LoVo cells

The proliferation of LoVo cells after P2RY10 silencing was assessed. The CCK-8 assay displayed no significant difference (*P* > 0.05) in cell viability between si-NC and si-P2RY10 cells after 24 h of culturing (**Figure 4A**). After culturing for 48 and 72 h, the viability of si-P2RY10-transfected LoVo cells was significantly lower than that of si-NC cells (*P* < 0.05; **Figure 4A**). Additionally, the si-NC and si-P2RY10 cells generated 451 ± 41 and 343 ± 47.1 colonies, respectively, indicating that si-P2RY10 transfection inhibited colony formation by LoVo cells (**Figure 4B**). Compared to si-NC cells (613 ± 36.4), the number of transmigrating si-P2RY10 cells (376 ± 92.4) was significantly decreased (*P* < 0.05; **Figure 4C**). The scratch-wound assay also revealed that the migration of LoVo cells was inhibited by P2RY10 silencing (**Figure 4D**). Collectively, these results suggested that P2RY10 silencing, similar to TRG-AS1 knockdown, significantly suppressed the viability, colony formation, and migration of LoVo cells.

**Figure 4.**
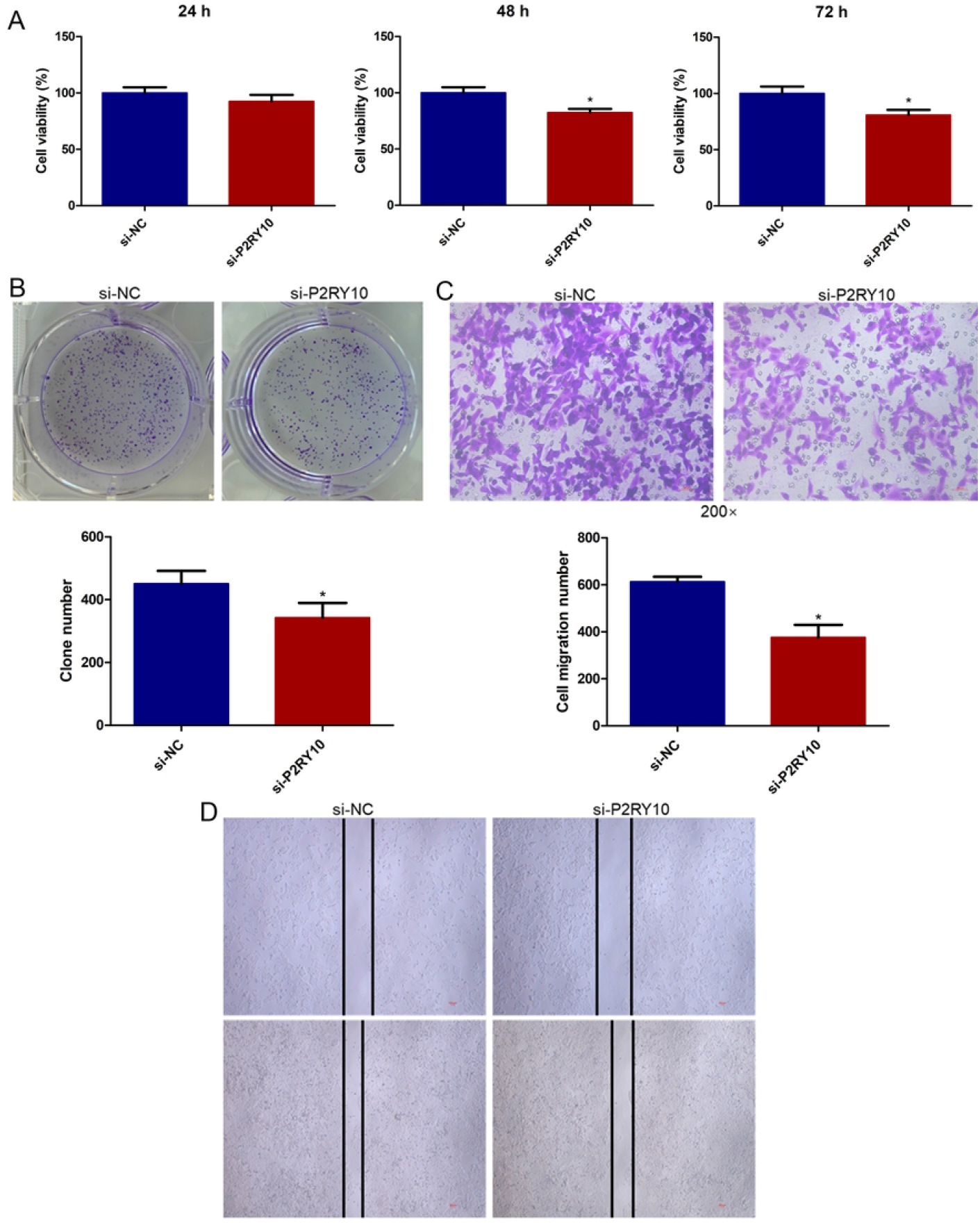
Effects of P2RY10 on LoVo cell proliferation. (**A**) Viability of LoVo cells after transfection with si-P2RY10 and cultured for 24, 48, and 72 h measured using the cell counting kit-8. (**B**) Transfected LoVo cell colony numbers by colony formation assay. Migration of transfected LoVo cells based on (**C**) Transwell® assay and (**D**) scratch test. * P < 0.05 vs. si-NC.

### 3.6 The role of GNA13 in the expression of P2RY10 and GNA13 in LoVo cells

P2RY10 has been demonstrated to activate GNA12/13 signaling, which is crucial for the proliferation of tumor cells [18, 17]. To further investigate the downstream pathways of P2RY10, LoVo cells overexpressing GNA13 were generated. The expression of GNA13 in the NC and OE-GNA13 cells was 1.54 ± 0.12 and 5932.10 ± 559.05, respectively; thus, GNA13 expression was approximately 6000 times higher in the OE-GNA13 group than that in the NC group (**Figure 5A**), indicating that LoVo cells overexpressing GNA13 were successfully generated and could be used for further study.

**Figure 5.**
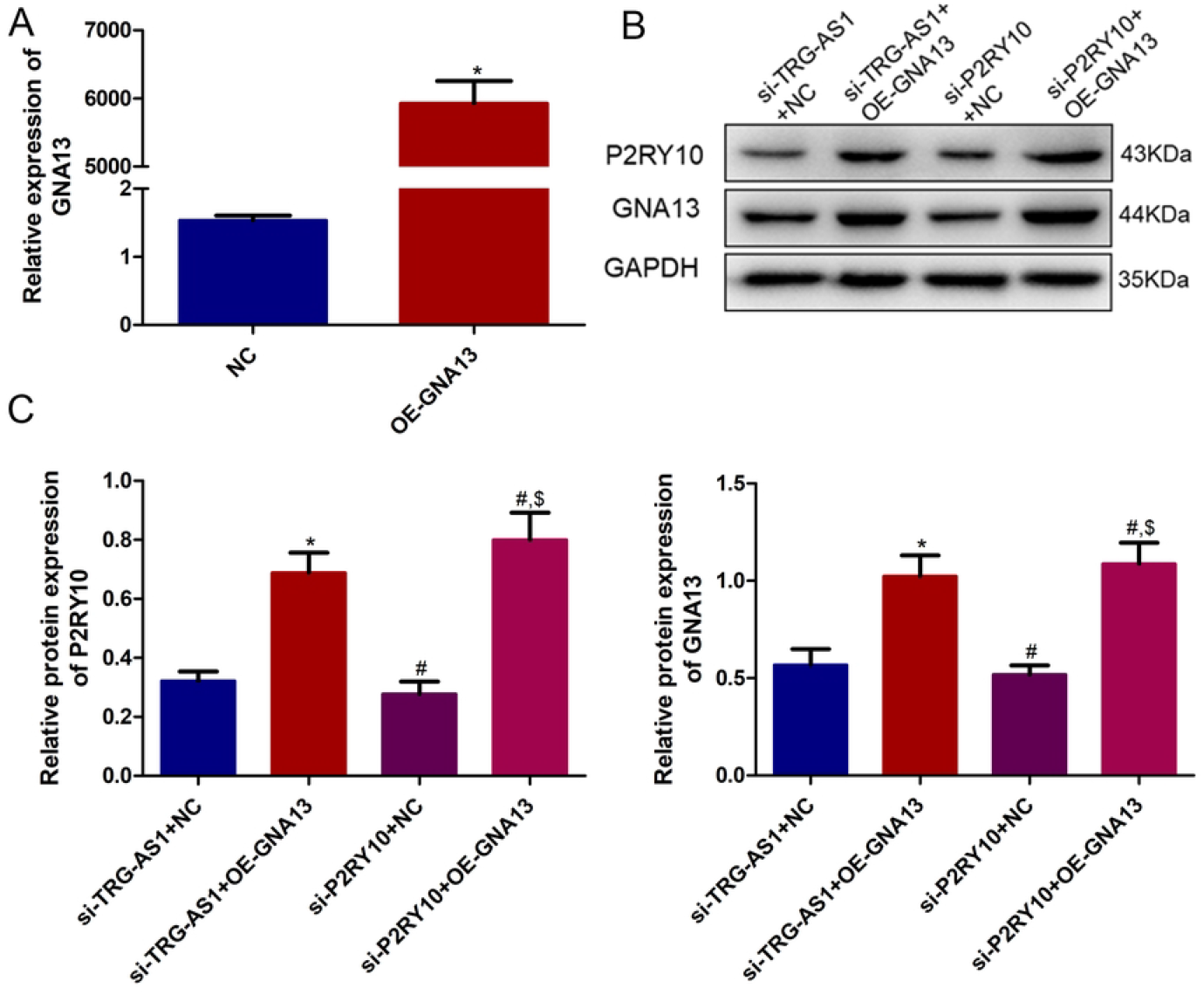
Effects of GNA13 on the expression of P2RY10 and GNA13 in LoVo cells. (**A**) LoVo cell transfection efficiency with OE-GNA13 plasmids. (**B**) Western blot of P2RY10 and GNA13 protein expression in transfected LoVo cells. * P < 0.05 vs. NC or si-TRG-AS1 + NC; # P < 0.05 vs. si-TRG-AS1 + OE-GNA13; $ P < 0.05 vs. si-P2RY10 + NC.

Subsequently, the P2RY10 and GNA13 protein levels were determined using western blotting. No significant differences (*P* > 0.05) were observed in the protein expression of P2RY10 and GNA13 between the si-TRG-AS1 + NC and the si-P2RY10 + NC cells (**Figures 5B, C**). In TRG-AS1 knockdown LoVo cells, GNA13 overexpression significantly upregulated P2RY10 and GNA13 protein levels relative to their corresponding NC cells (*P* < 0.05; **Figures 5B, C**). In P2RY10-silenced LoVo cells, the protein expression of P2RY10 and GNA13 was upregulated by GNA13 overexpression compared with the NC cells (*P* < 0.05; **Figures 5B, C**). In addition, P2RY10 and GNA13 protein levels were significantly higher in the si-P2RY10 + OE-GNA13 cells compared with the si-TRG-AS1 + OE-GNA13 cells (*P* < 0.05; **Figures 5B, C**).

### 3.7 Role of GNA13 in the proliferation of LoVo cells

When cultured for 24 h, cell viability was not significantly different among the different cells (*P* > 0.05, **Figure 6A**). When LoVo cells with TRG-AS1 knockdown or P2RY10 silencing were cultured for 48 and 72 h, GNA13 overexpression significantly (*P* < 0.05) enhanced the viability of LoVo cells relative to that of the corresponding NC cells (**Figure 6A**). The colony-forming ability of cells overexpressing GNA13 was significantly higher (*P* < 0.05) than that of the corresponding NC cells (**Figure 6B**). Finally, Transwell® and scratch-wound assays showed that GNA13 overexpression increased the migration of LoVo cells with TRG-AS1 knockdown or P2RY10 silencing (**Figures 6C, D**). Taken together, the overexpression of GNA13 reversed the reduction in viability, colony formation, and migration of TRG-AS1 knockdown or P2RY10-silenced LoVo cells.

**Figure 6.**
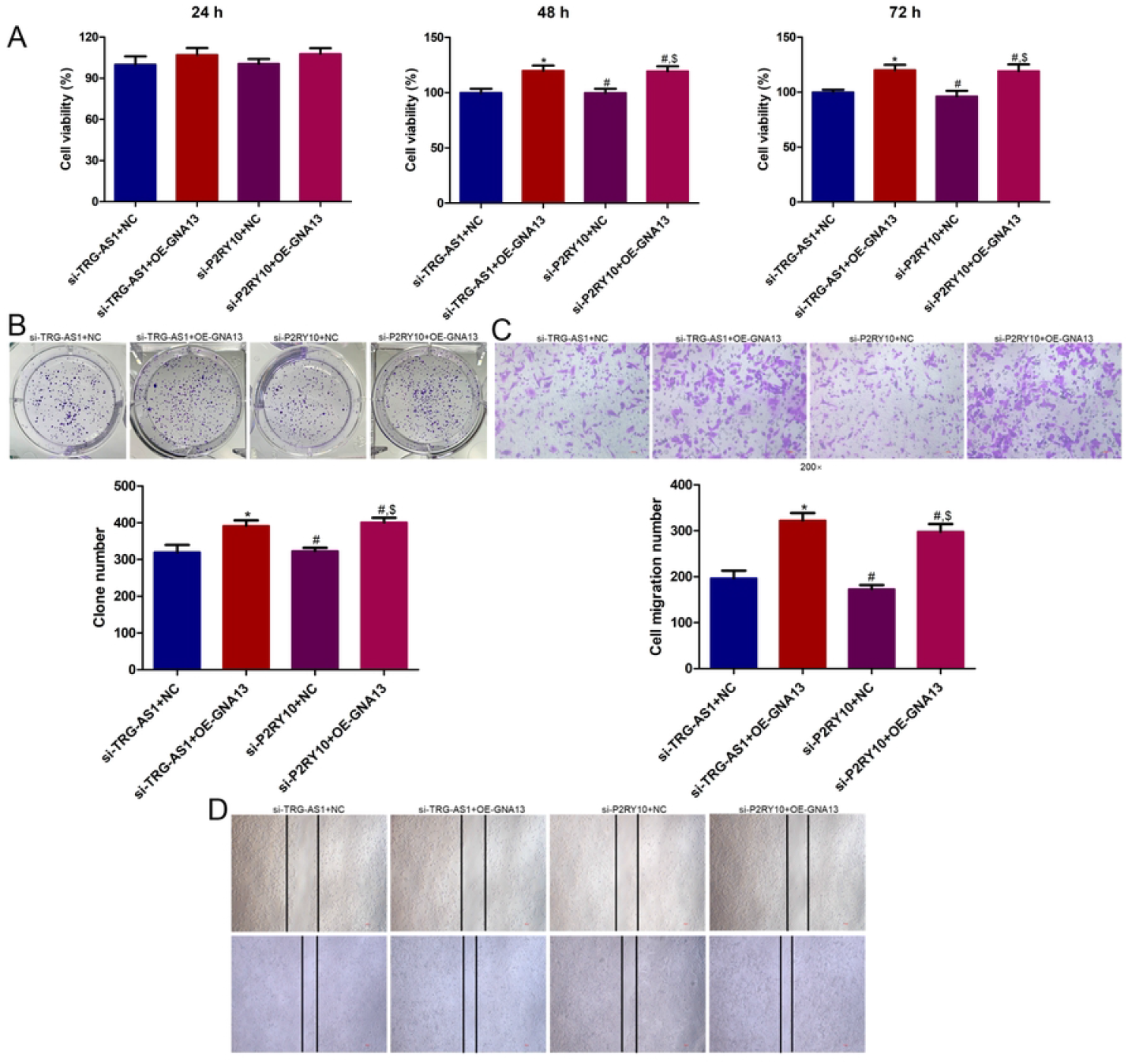
Effects of GNA13 on LoVo cell proliferation. LoVo cells were transfected with plasmids for TRG-AS1 knockdown or P2RY10 silencing and overexpression of GNA13. (**A**) Viability of transfected LoVo cells cultured for 24, 48, and 72 h measured using the cell counting kit-8. (**B**) Transfected LoVo cell colony numbers based on colony formation assay. Transfected LoVo cell migration based on (**C**) Transwell® assay and (**D**) scratch test. * P < 0.05 vs. si-TRG-AS1 + NC; # P < 0.05 vs. si-TRG-AS1 + OE-GNA13; $ P < 0.05 vs. si-P2RY10 + NC.

### 3.8 Effects of TRG-AS1 and P2RY10 on tumor growth

To further explore the roles of TRG-AS1 and P2RY10 in CRC *in vivo*, nude mice were subcutaneously injected with different cell suspensions, and tumor volumes were monitored weekly. We found that tumor volumes in the si-TRG-AS1 and si-P2RY10 groups were smaller at the end of the experiment (**Figure 7A**). Over time, the tumor volumes in the si-NC group were significantly larger (*P* < 0.05) than those in the si-TRG-AS1 and si-P2RY10 groups (**Figure 7B**). Tumor volumes in the si-TRG-AS1 group (*P* > 0.05) were similar to those in the si-P2RY10 group (**Figure 7B**). Additionally, HE staining of the colon tumor tissue morphology showed that a large number of tumor cells infiltrated the colon tissues in the si-NC group, whereas there was very limited invasion by tumor cells in the si-TRG-AS1 and si-P2RY10 groups (**Figure 7C**).

**Figure 7.**
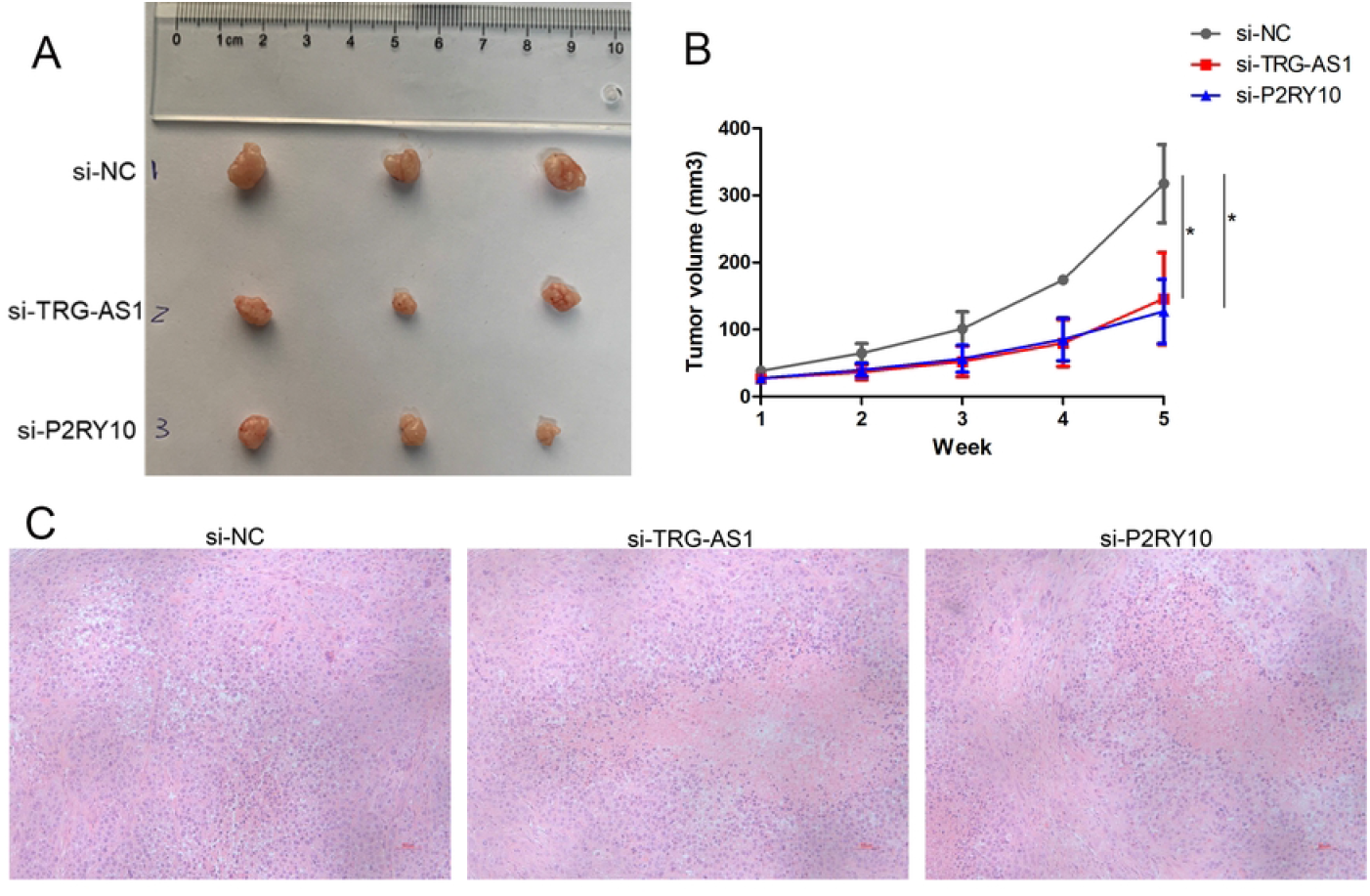
Effects of TRG-AS1 and P2RY10 on tumor growth *in vivo*. (**A**) Tumor volumes in the different groups at the end of the experiment. (**B**) Tumor volume changes in the different groups over time. (**C**) Morphology of the tumor tissues in the different groups observed by hematoxylin and eosin staining. * P < 0.05. No.

## 4 Discussion

CRC remains the leading cause of cancer-related deaths worldwide, with less than half of the cases diagnosed when the cancer is locally advanced, and is considered a global health problem [20]. Our previous bioinformatics analysis indicated that lncRNA TRG-AS1 and its co-expressed gene P2RY10 are important for CRC occurrence and development, according to the degree of correlation between this co-expression network and the Pearson correlation coefficient [13]. The current research further verified the specific roles of TRG-AS1 and P2RY10 in CRC using both *in vitro* and *in vivo* experiments. TRG-AS1 knockdown and P2RY10 silencing suppressed LoVo cell viability, colony formation, and migration. GNA13 overexpression reversed changes in LoVo cells induced by TRG-AS1 knockdown or P2RY10 silencing.

LncRNAs act as different functional molecules (signals and scaffolds) in various cancer-related cellular processes by interacting with co-activators of RNA-binding proteins and transcription factors or by altering the primary promoters of their targets [21]. Wang et al. [22] demonstrated that the lncRNA LINRIS was upregulated in the tissues of CRC patients with poor OS, and suppression of LINRIS resulted in impaired growth of CRC cell lines and inhibited tumor growth *in vivo* via downregulation of IGF2BP2 and reduction of MYC-mediated glycolysis by CRC cells, thus representing a promising target for CRC therapy. Another study showed that lncRNA NEAT1 levels were clearly increased in CRC tissues relative to control tissues and that it could promote CRC metastasis and progression via Wnt/β-catenin signaling by direct interaction with DDX5 [23]. Therefore, lncRNA expression is closely associated with cancer pathogenesis and development. The lncRNA TRG-AS1 was found to be more abundant in hepatocellular carcinoma (HCC) cells, and its knockdown inhibited HCC cell propagation and EMT by targeting the miR-4500/BACH1 axis [24]. Xie et al. [25] reported that TRG-AS1 was significantly associated with a poor prognosis and was upregulated in glioblastoma tissues and cells. In addition, by acting as a competing endogenous RNA (ceRNA) for miR-877-5p, TRG-AS1 mediated *SUZ12* expression and promoted glioblastoma cell proliferation, thus representing a novel therapeutic target for glioblastoma. In CRC, *P2RY10* is co-expressed with lncRNA TRG-AS1 and is associated with chemotaxis through eosinophil degranulation, which may be a potential target for cancer treatment [26]. P2RY10 expression is increased in synovial tissues and peripheral blood in patients with rheumatoid arthritis and coronary artery disease [27], and the LysoPS-P2Y10 axis has been shown to inhibit TNF-α production in mouse microglia and activate eosinophilic degranulation [17]. A previous study demonstrated that P2RY10 was related to the tumor microenvironment and was a crucial prognostic factor in cutaneous melanoma [28]. Another study showed that P2RY10 was an immune/matrix-related gene that predicted the survival rate of patients with renal cell carcinoma [29]. Our study also showed that TRG-AS1 knockdown downregulated P2RY10 expression, and that P2RY10 silencing downregulated TRG-AS1 expression. Taken together, we speculate that the suppression of TRG-AS1 and P2RY10 may reduce CRC progression by inhibiting the proliferation of CRC cells and that TRG-AS1 and P2RY10 may interact with each other.

P2RY10 activates GNA12/13 signaling and triggers the Rho/Rho-associated kinase pathway [30]. Additionally, the increased expression and enhanced signal transduction of *GNA12* and *GNA13* has been associated with tumor progression and tumorigenesis in many types of cancers [31]. Our study showed that TRG-AS1 knockdown and P2RY10 silencing decreased GNA12 and GNA13 expression. Therefore, we established GNA13-overexpressing cells to further explore the relationship between TRG-AS1, P2RY10, and their downstream functions. GNA13 overexpression reversed the reduction in P2RY10 expression and cell proliferation caused by TRG-AS1 knockdown or P2RY10 silencing in LoVo cells. GNA12 is a heterotrimeric G protein α subunit with carcinogenic potential. GNA12 activation *in vitro* and *in vivo* promotes the invasion of breast and prostate cancer cells, and its expression is upregulated in many tumors, especially in metastatic tissues [32]. Ha et al. [33] found that GNA12 could drive ovarian cancer progression by upregulating tumor-promoting networks with *VEGFA*, insulin-like growth factor 1, *STAT3, AKT1, BCL2L1*, growth hormone-releasing hormone, and *TGFB1* as key hubs/nodes. GNA13 is an essential factor for EMT in the process of CRC metastasis [34] and regulates angiogenesis by inducing VEGFR2 expression [35]. Zhang et al. [36] reported that GNA13 was significantly overexpressed in gastric cancer (GC) tissues and was significantly related to positive GC progression and adverse survival outcomes. Silencing of GNA13 expression *in vitro* and *in vivo* significantly suppressed the proliferation and tumorigenicity of GC cells, suggesting that GNA13 may be a target for GC therapy in humans. Another study demonstrated that GNA13 expression was essential for the invasion and migration of prostate cancer cells and was post-transcriptionally regulated by miR-182 and miR-200 [37]. Together with our findings, these reports suggest that TRG-AS1 knockdown represses CRC cell proliferation by regulating P2RY10 and GNA13 expression, thereby controlling CRC occurrence and development.

In conclusion, both TRG-AS1 and P2RY10 were upregulated in CRC cells, and the suppression of TRG-AS1 and P2RY10 reduced CRC progression by inhibiting the growth of CRC cells. Possible mechanisms included expression of P2RY10 and GNA13. These findings provide a theoretical foundation for the management of CRC using TRG-AS1/P2RY10/GNA13 as potential therapeutic targets.

## 6 Author Contributions

DLS, LZ and HZ conceived and designed the experiments. LZ, BYL, QZ, YJL and LHD performed the experiments. QTZ, BYL, QZ and LHD analyzed the data. LZ and HZ wrote the article. All authors read and approved the final manuscript.

## 8 Acknowledgments

We are thankful to our laboratory colleagues for their assistance in our research.

## 9 Ethics declarations

Human specimen collection was approved by the Ethics Committee of Wujin Affiliated Hospital of Jiangsu University. Written informed consent was obtained from each patient according to the policies of the committee. The animal protocol was approved by the Institutional Animal Care and Use Committee of Nanjing Medical University (Protocol Number NJMU08-092).

Data will be made available on request.

